# Evaluation of the Potential of Two Common Pacific Coast Macroalgae for Mitigating Methane Emissions from Ruminants

**DOI:** 10.1101/434480

**Authors:** Charles G. Brooke, Breanna M. Roque, Negeen Najafi, Maria Gonzalez, Abigail Pfefferlen, Vannesa DeAnda, David W. Ginsburg, Maddelyn C. Harden, Sergey V. Nuzhdin, Joan King Salwen, Ermias Kebreab, Matthias Hess

## Abstract

With increasing interest in feed based methane mitigation strategies, fueled by local legal directives aimed at methane production from the agricultural sector in California, identifying local sources of biological feed additives will be critical in keeping the implementation of these strategies affordable. In a recent study, the red alga *Asparagopsis taxiformis* stood out as the most effective species of seaweed to reduce methane production from enteric fermentation. Due to the potential differences in effectiveness based on the location from where *A. taxiformis* is collected and the financial burden of collection and transport, we tested the potential of *A. taxiformis*, as well as the brown seaweed *Zonaria farlowii* collected in the nearshore waters off Santa Catalina Island, CA, USA, for their ability to mitigate methane production during *in-vitro* rumen fermentation. At a dose rate of 5% dry matter (DM), *A. taxiformis* reduced methane production by 74% (*p* ≤ 0.01) and *Z. farlowii* reduced methane production by 11% (*p* ≤ 0.04) after 48 hours and 24 hours of *in-vitro* rumen fermentation respectively. The methane reducing effect of *A. taxiformis* and *Z. farlowii* described here make these local macroalgae promising candidates for biotic methane mitigation strategies in the largest milk producing state in the US. To determine their real potential as methane mitigating feed supplements in the dairy industry, their effect *in-vivo* requires investigation.

## 1. Introduction

Methane (CH_4_) accounts for more than 10% of the greenhouse gas (GHG) emissions from the US (Myhre, 2013) and enteric fermentation from ruminant animals accounts for approximately 25% of the total anthropogenically produced methane (NASEM, 2018). Thus, efficient strategies to lower enteric CH_4_ production could result in a significantly reduced carbon footprint from agriculture and animal production more specifically.

*In-vitro* studies have demonstrated that some brown and red macroalgae can inhibit microbial methanogenesis (Machado, 2014) and they have been suggested as feed supplements to reduce methanogenesis during enteric fermentation (Machado, 2016; Dubois, 2013; Wang, 2008). In addition to its methane reducing affect, utilization of these macroalgae could promote higher growth rates and feed conversion efficiencies in ruminants via the potential net energy yield from the redistribution of energy from the microbial methanogenesis pathway, into more favorable pathways (i.e volatile fatty acids) (Hansen, 2003; Marín, 2009). Therefore, macroalgae feed supplementation may be an effective strategy to simultaneously improve profitability and sustainability of beef and dairy operations.

In a recent study (Machado et al. 2014), the red alga *Asparagopsis taxiformis* stood out as the most effective species of seaweed to reduce methane production. In this work, the effect of a large variety of macroalgal species including freshwater, green, red and brown algae on CH_4_ production during *in-vitro* incubation were compared and the obtained results showed that *A. taxiformis* amendment yielded the most significant reduction (~98.9%) of CH_4_ production.

A major barrier to the implementation of an *A. taxiformis* based methane mitigation strategy is the availability of the seaweed, which has led to the exploration of alternative seaweed species. Previous investigations have collected *A. taxiformis* during diving excursions off the coast of Australia. Due to the potential differences in effectiveness based on the location and growing conditions from which the seaweed is collected and the financial burden of transport, we tested the potential of two different species of subtidal macroalgae (*A. taxiformis* and the brown alga, *Zonaria farlowii*) from Southern California for their ability to mitigate methane production during *in-vitro* rumen fermentation.

## 2. Materials and Methods

### 2.1 Experimental Design

To determine the effect of two locally sourced macroalgae species on methane production during in-vitro rumen fermentation, *Asparagopsis taxiformis* and *Zonaria farlowii* were supplemented to an *in-vitro* gas production system at a dose rate of 5% DM. Rumen fluid was diluted 3-fold with artificial saliva buffer (Oeztuerk et al., 2015). After homogenization, 200 ml of the mixture was allocated to 300 ml vessels fitted with Ankom head units (Ankom Technology RF Gas Production System, Macedon, NY, USA). Each vessel recieved 2 g of rumen solids, and 2 g of a basic ration (Super basic ration — SBR, Table 1) commonly used in the dairy industry in California. Rumen solids and SBR were sealed in separate Ankom feed bags and seaweed was included in the respective SBR feed bags (Ankom, Macedon, NY). Vessels were placed in a shaking water bath (39°C) and incubated while mixed at 40 rpm. Foil gas bags (Restek, USA) were connected to the Ankom head units to collect gas at 24 and 48 hours respectively.

### 2.2 Pacific Coast Seaweed Collection and Preparation

*Asparagopsis taxiformis* and *Z. farlowii* were collected from Little Fisherman’s Cove on the leeward side of Santa Catalina Island, ~35 km off the coast of Southern California, USA (Figure 1). The seaweed was shipped on ice to the University of California, Davis, where it was dried at 55°C for 72 hours and ground through a 2 mm Wiley Mill (Thomas Scientific, Swedesboro, NJ).

**Figure 1.**
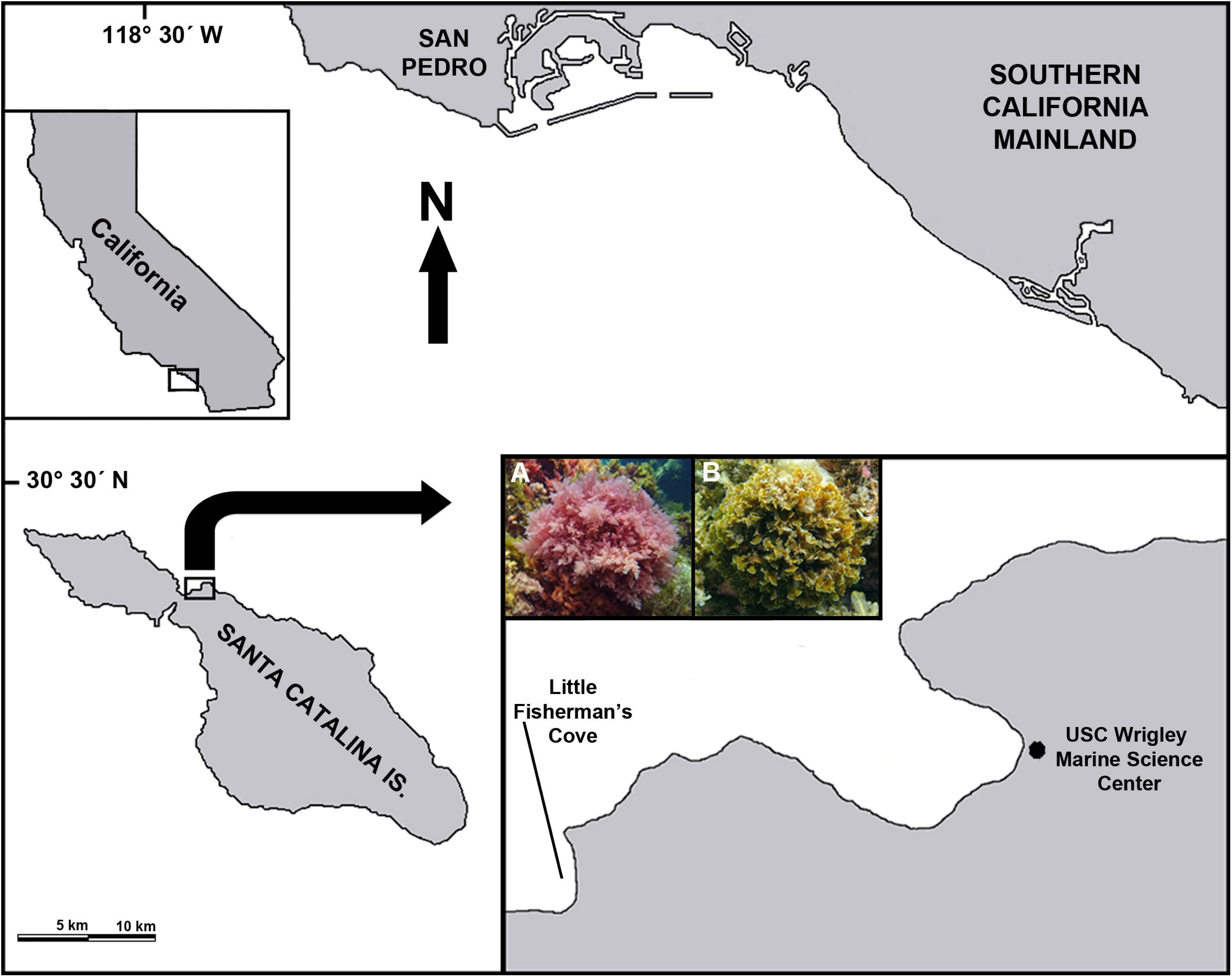
Map showing the location of Santa Catalina Island relative to the Southern California mainland. Inset: The red alga *Asparagopsis taxiformis* (A) and the brown alga *Zonaria farlowii* (B) were collected (2-5 m depth) in Little Fisherman’s Cove, located ~0.6 km from the USC Wrigley Marine Science Center.

### 2.3 Rumen Fluid Collection

All animal procedures were performed in accordance with the Institution of Animal Care and Use Committee (IACUC) at University of California, Davis under protocol number 19263. Rumen content was collected from a rumen fistulated Holstein cow, housed at the UC Davis Dairy Research and Teaching Facility Unit. The rumen fluid donor was fed a dry cow total mixed ration (50% wheat hay, 25% alfalfa hay/manger cleanings, 21.4% almond hulls, and 3.6% mineral pellet, Table 1). Two liters of rumen fluid and 30 g of rumen solids were collected 90 min after morning feeding. Rumen content was collected via transphonation using a perforated PVC pipe, 500 mL syringe, and Tygon tubing (Saint-Gobain North America, PA, USA). Fluid was strained through a colander and 4 layers of cheesecloth into a 4 L pre-warmed, vacuum insulated container and transported to the laboratory.

### 2.4 Greenhouse Gas Analysis

Methane and CO_2_ were measured from gas bags using an SRI Gas Chromatograph (8610C, SRI, Torrance, CA) fitted with a 3’x1/8” stainless steel Haysep D column and a flame ionization detector (FID) with methanizer. The oven temperature was held at 90°C for 5 minutes. Carrier gas was high purity hydrogen at a flow rate of 30 ml/min. The FID was held at 300°C. A 1 mL sample was injected directly onto the column. Calibration curves were developed with Airgas certified CH_4_ and CO_2_ standard (Airgas, USA).

### 2.5 Statistical Analysis

Differences in CH_4_ and CO_2_ production were determined using unpaired parametric t-tests with Welch’s correction conducted in Graphpad Prism 7 (Graphpad software Inc, La Jolla, CA). Significant differences among treatments were declared at *p* ≤ 0.05.

## 3. Results

### 3.1 Gas production profile of *in vitro* fermentation of rumen fluid amended with 5% *A. taxiformis*

At a dose rate of 5% DM, *A. taxiformis* reduced methane production by 74% after 48 hours of *in-vitro* rumen fermentation (*p* ≤ 0.01, Figure 2B) and daily methane production remained nearly identical in the presence of *A. taxiformis* on both days (7.1±1.9 ml (g DM)^-1 and 6.6±2.5 ml (g DM)^-1 after 24 and 48 hours respectively). Methane production in the control vessels increased by 76% after 48 hours of incubation (20.3±11 ml (g DM)^-1 and 35.5±8.5 ml (g DM)^-1 at 24 and 48 hours respectively).

**Figure 2.**
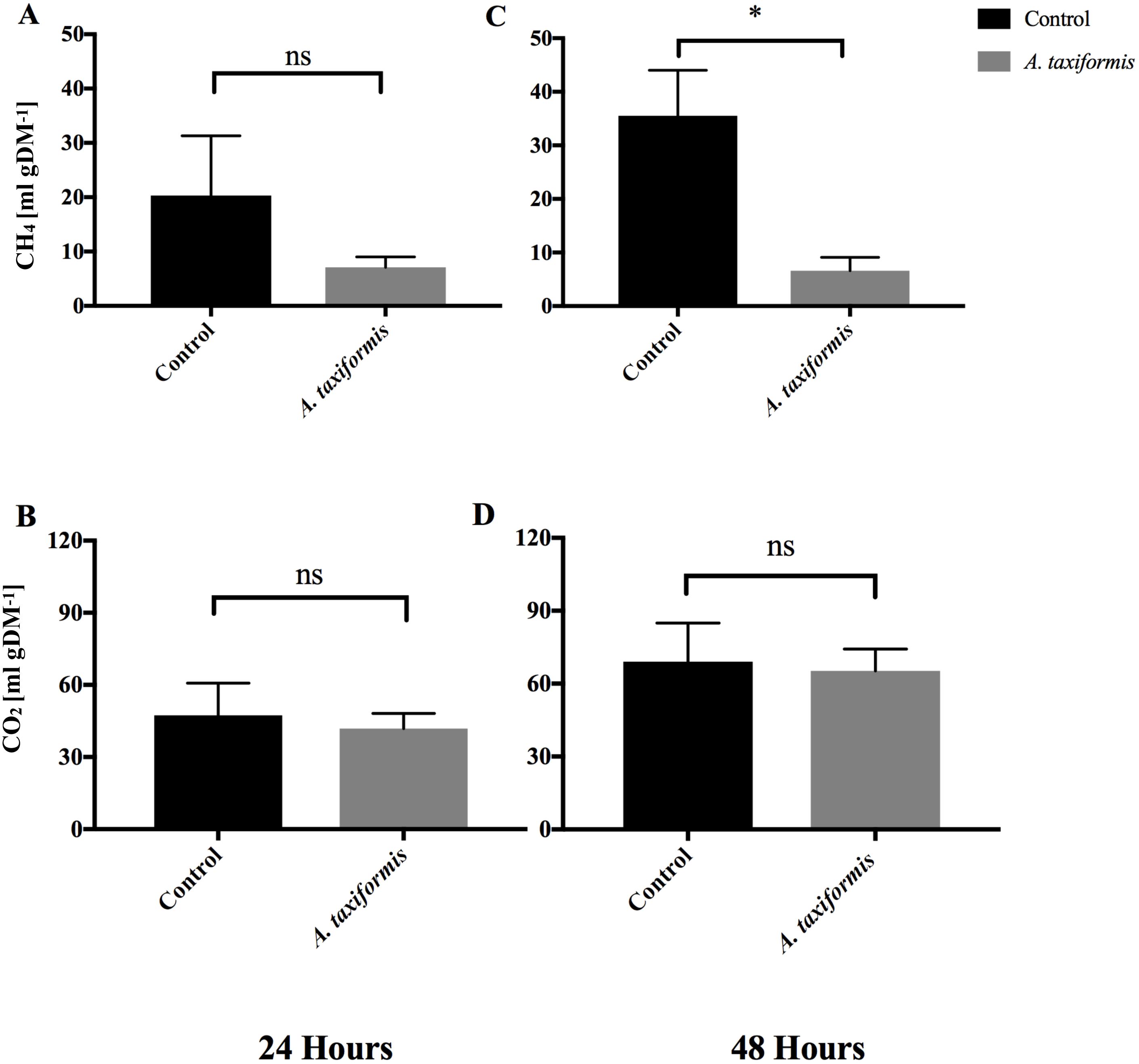
Methane and CO_2_ production during *in-vitro* fermentation of rumen fluid amended with *A. taxiformis*. Production of CH_4_ [ml (g DM)^-1] and CO_2_ [ml (g DM)^-1] from vessels without (n=4) and with 5% (n=4) *A. taxiformis* as additive. Methane and CO_2_ were measured at 24 h (**A** & **B** respectively) and 48 h (**C** & **D** respectively). “*“ indicate significant difference (*p* value ≤ 0.05), “ns” indicates not significant. Error bars represent the standard error from the mean.

While methane production varied with 5% DM inclusion of *A. taxiformis,* CO_2_ production remained similar between treatment (41.9±6.2 ml (g DM)^-1 and 65.23±9.1 ml (g DM)^-1 at 24 and 48 hours respectively) and control vessels (47.4±13.4 ml (g DM)^-1 and 69.0±15.9 ml (g DM)^-1 at 24 and 48 hours respectively).

### 3.2 Gas production profile of *in vitro* fermentation of rumen fluid amended with 5% *Z. farlowii*

At a dose rate of 5% DM, *Z. farlowii* reduced methane production by 11% after 24 hours of *in vitro* rumen fermentation (*p* ≤ 0.04, Figure 3A). Daily methane production decreased slightly at 48 hours compared to 24 hours of incubation for both the control and treatment vessels (Control = 62.5±3.3 ml (g DM)^-1 and 51.4 ±2.9 ml (g DM)^-1 CH_4_, at 24 and 48 hours respectively; treatment = 55.3 ±2.7 and 45.9 ±3.7 ml (g DM)^-1 CH_4_, at 24 and 48 hours respectively).

**Figure 3.**
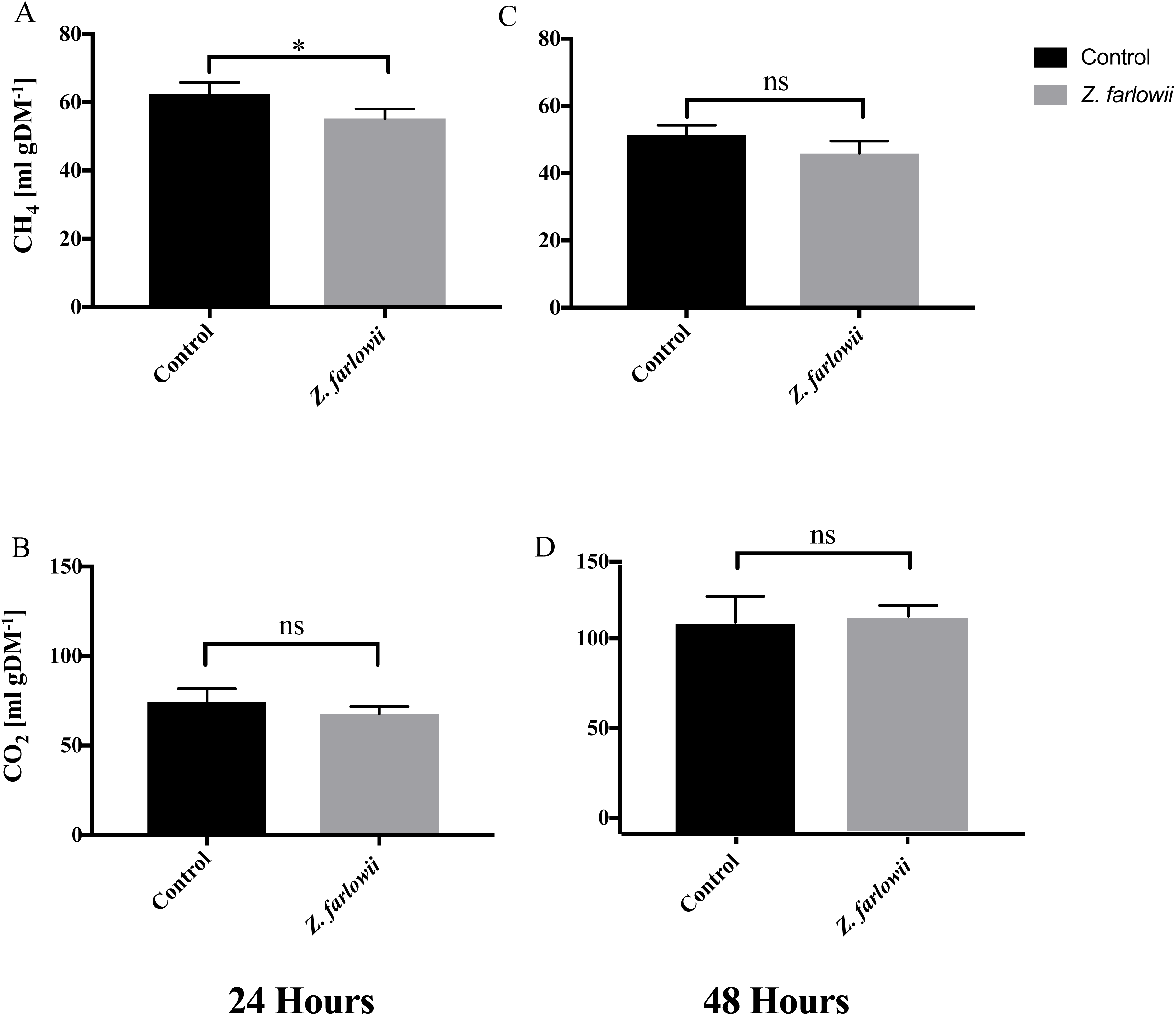
Methane and CO_2_ production during *in-vitro* fermentation of rumen fluid amended with *Z. farlowii*. Production of CH_4_ [ml (g DM)^-1] and CO_2_ [ml (g DM)^-1] from vessels without (n=3) and with 5% (n=3) *Z. farlowii* as additive. Methane and CO_2_ were measured at 24 h (**A** & **B** respectively) and 48 h (**C** & **D** respectively). “*“ indicate significant difference (*p* value ≤ 0.05),”ns” indicates not significant. Error bars represent the standard error from the mean.

While methane production decreased slightly for all vessels at 48 hours, CO_2_ production nearly doubled (Control = 74.1±7.7 ml (g DM)^-1 and 117.9 ±14.6 ml (g DM)^-1 CO_2_, at 24 and 48 hours respectively; treatment = 67.6±4.1 ml (g DM)^-1 and 114.2±6.0 ml (g DM)^-1 CO_2_, at 24 and 48 hours respectively). Carbon dioxide production from vessels amended with 5% DM of *Z. farlowii* did not differ from the control vessels at 24 or 48 hours (*p* ≤ 0.27 and *p* ≤ 0.70 respectively).

## 4. Discussion

With increasing interest in feed-based biotic methane mitigation strategies fueled by legal directives aimed at reducing methane production from the agricultural sector, identification of local biotic feed-supplements will be critical to render large-scale methane mitigation strategies economical.

The data presented here suggest that subtidal macroalgae from Santa Catalina Island, Southern California reduced the *in-vitro* production of CH_4_ when added to rumen content from California dairy cattle, suggesting that California seaweed might represent a viable option for use in feed based methane mitigation strategies. In addition to demonstrating the potential of the local *A. taxiformis* for methane mitigation during enteric fermentation, we also demonstrated significant methane reduction in the brown alga *Z. farlowii,* a species of seaweed commonly found along the Southern California Bight, without obvious impact on CO_2_ production (Figures 2 and 3, panels A and B).

The effectiveness of a macroalgae in reducing methane production during rumen incubation has been linked to the concentration of halogenated bioactives including bromoform and di-bromochloromethane (Machado, 2016). However, in contrast to *A. taxiformis,* which has been shown to produce several halomethane compounds, *Z. farlowii* amendment only reduced methane on a short time scale. These findings suggest that either the bioactives in *Z. farlowii* are more bioavailable but less effective or concentrated, or methane reduction is occurring via a different compound or a different mode of action. Previous studies have identified multiple phenolic lipids produced by *Z. farlowii* from Southern California waters as possessing antimicrobial activity (Gerwick and Fenical, 1981). However, the reduction of methane in vessels amended with *Z. farlowii* was modest compared to those amended with *A. taxiformis*. *Zonaria farlowii* is commonly found along the Southern California Bight, which makes it a potential candidate for non-terrestrial farming operations along the Southern California Coast. A more in-depth nutrient analysis of *Z. farlowii* along with *in-vitro* assays will be essential to help determine its value for future methane mitigation strategies and to determine its potential for use in dairy operations.

## 5. Conclusion

*Asparagopsis taxiformis* and *Z. farlowii* collected off Santa Catalina Island were evaluated for their ability to reduce methane production from dairy cattle fed a mixed ration widely utilized in California. The methane reducing effect of the *A. taxiformis* and *Z. farlowii* described in this study makes these macroalgae promising candidates for biotic methane mitigation strategies in the largest milk producing state in the US. With expected growth in livestock production, it is necessary to investigate and confirm the effect of these macroalgae *in-vivo,* in order to ensure that farmers have sufficient incentive to implement such strategies.

